# Melittin-induced alterations in morphology and deformability of human red blood cells using quantitative phase imaging techniques

**DOI:** 10.1101/091991

**Authors:** Joonseok Hur, Kyoohyun Kim, SangYun Lee, HyunJoo Park, YongKeun Park

## Abstract

Here, the actions of melittin, the active molecule of apitoxin or bee venom, were investigated on human red blood cells (RBCs) using quantitative phase imaging techniques. High-resolution realtime 3-D refractive index (RI) measurements and dynamic 2-D phase images of individual melittin-bound RBCs enabled in-depth examination of melittin-induced biophysical alterations of the cells. From the measurements, morphological, biochemical, and mechanical alterations of the RBCs were analyzed quantitatively. Furthermore, leakage of haemoglobin (Hb) inside the RBCs at high melittin concentration was also investigated.

## Introduction

Melittin, the active molecule of apitoxin or bee venom, is a transmembrane protein, and it forms small pores on the cell membrane^1^. Once melittin binds to the lipid membranes of cells, toroid-shaped pores are formed and enable the leakage of molecules with the size of tens of kDa. It results in changes in the permeability of the cell membrane depending on the melittin concentration. As a pore-forming protein, melittin has been extensively studied from diverse aspects including its structure, binding mechanisms, and pore-forming processes^2–10^. Because of its intriguing interaction with lipid membranes and its pore-forming capability, melittin has potential to be used for various applications including antimicrobial, cell-selective attack, and translocation of materials by changing the membrane permeability^11–13^.

Despite the potential uses of melittin, the cellular action of melittin has not been fully investigated mostly due to technical limitations. Previous studies have focused on observing melittin-induced cell lysis rather than measuring characteristic biophysical alterations of cells during or before lysis. This is mainly due to the lack of quantitative and high-resolution techniques for visualizing individual cells. For instance, bright-field microscopy, phase contrast microscopy, and differential interference contrast microscopy provide qualitative information about cells^12,14–16.^ Some quantitative studies have been done using flow-based imaging techniques; however, these techniques are subject to low precision as well as difficulties in investigating the dynamics of cells^15,17,18^. Dynamic alterations of live cells cannot be measured with electron microscopy either^15,17^, and fluorescent imaging enables only selective measurements of labelled parts^12,15,17^.

Recent developments in quantitative phase imaging (QPI) techniques have enabled quantitative, high-resolution, 3-D, real-time, label-free, and non-invasive measurements of individual cells and thus have provided the capability of overcoming conventional experimental limitations^19–21^. The 3-D morphologies and internal configuration of cells with associated biophysical parameters including cell volumes, surface areas, and dry masses can be measured with holographic optical tomography^22–26^. Additionally, 2-D dynamic phase maps can measure the rapid dynamics of a cell at a time resolution of milliseconds^27–30^. Furthermore, a recent improvement in holographical optical tomography enables 3-D measurements of cells with a time resolution of hundreds of milliseconds^31^, which was a sufficient resolution to measure melittin-induced alterations of cells (see Results).

Here, we present the measurements of alterations in the morphologies and deformability of live red blood cells (RBCs) induced by melittin. High-resolution 3-D refractive-index (RI) maps and 2-D dynamic phase shift maps of individual melittin-associated RBCs were measured by common-path diffraction optical tomography (cDOT). Among QPI techniques, cDOT has a sensitivity of a few nanometres and enables the simultaneous measurement of 3-D RI tomograms and 2-D dynamic phase images in the same setup with a common-path configuration^32^. Human RBCs were chosen as a model cell to study the membrane-acting biochemistry because they have a simple cellular structure and are suitable for optical measurements by optical contrast from cytoplasmic haemoglobin (Hb). Furthermore, the 2-D spectrin network underlying the plasma membrane of RBCs provides the unique mechanical properties and deformability of RBCs, which have been of interest in numerous studies^22,29,30,33–36^. It was observed that melittin induces characteristic morphological changes in RBCs called echinocytosis. The phase of echinocytosis was dependent on the concentration of melittin. Quantitative morphological parameters including cell volume, surface area, and sphericity were further investigated. The level of membrane fluctuation in the RBC decreased when melittin was introduced indicating a loss of deformability of the RBC. Real-time measurement of the melittin-induced changes in the RBCs showed the leakage of Hb with a constant morphology suggesting the formation of a stable pore of sufficient size with a high concentration of melittin. This study reports on the 3-D investigation of melittin-induced alterations in human RBCs at the single cell level and demonstrates systematic approaches to studying the cellular actions of membrane-active molecules using cDOT.

## Results

### 3-D RI tomograms of individual RBCs exposed to various melittin concentrations

To investigate how melittin alters the morphologies of RBCs as a transmembrane protein, the 3-D RI tomograms of individual RBCs were measured using cDOT with various concentrations of melittin from 50 to 500 nM as shown in Fig. 1A–D. The 3-D RI tomograms of the RBCs showed significant morphological changes as the melittin concentration was increased; RBCs underwent morphological transitions from discocytes to echinocytes and to spherocytes. Before treated with melittin, the RBCs exhibited the characteristic thin, doughnut-like shapes or discocyte (Fig. 1A).

**Fig. 1.**
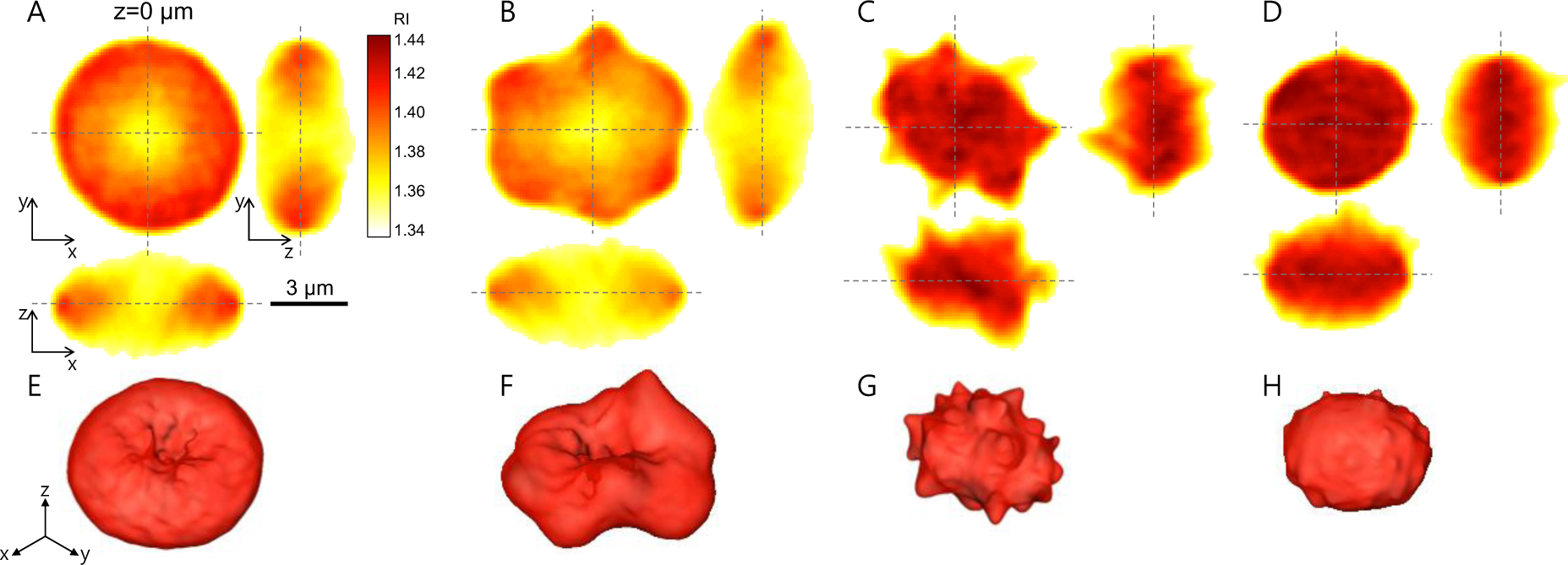
(A-D) Cross-sectional slices of reconstructed RI tomograms of RBCs at various melittin concentrations of 0 (A), 50 (B), 200 (C), and 500 (D) nM in *x-y* (left),*y-z* (right), and x-z (below) planes. (E-H) Corresponding rendered isosurfaces of the 3-D RI maps (n > 1.368).

As the concentration of the melittin solution increased, the RBCs started to undergo morphological alterations resulting in irregular membrane shapes. At a melittin concentration of 50 nM, RBC membranes exhibited bumpy structures while maintaining their dimple structure in the centre (Fig. 1B). When the concentration of melittin was 200 nM, the RBCs lost their dimple shapes, and specular structures emerged in the membranes, or they became echinocytes (Fig. 1C). At a concentration of 500 nM, the RBCs became spherical shapes in which the specular structure disappeared. These morphological transitions can also be seen in the isosurface images of the reconstructed RI tomograms (Figs. 1E–H), which were consistent with previous reports in which bright-field microscopy was used^14,16^.

For quantitative analysis, morphological parameters including cell volume, surface area, and sphericity were extracted from the measured 3-D RI tomograms (see Methods). As shown in Fig. 2, the volumes of the RBCs did not significantly decrease up to melittin concentrations of 150 nM. At concentrations of 200 nM and above, cell volumes decreased by 12%. The mean values of the RBC cell volumes for melittin concentrations of 0, 50, 100, 150, 200, 300 and 500 nM were 90.5 ± 10.4 (N = 35), 90.3 ± 12.24 (N = 34), 90.8 ± 12.24 (N = 33), 88.0 ± 11.64 (N = 33), 79.6 ± 11.14 (N = 33), 79.2 ± 9.54 (N = 36), and 77.6 ± 8.74 fL (N = 32), respectively.

**Fig. 2.**
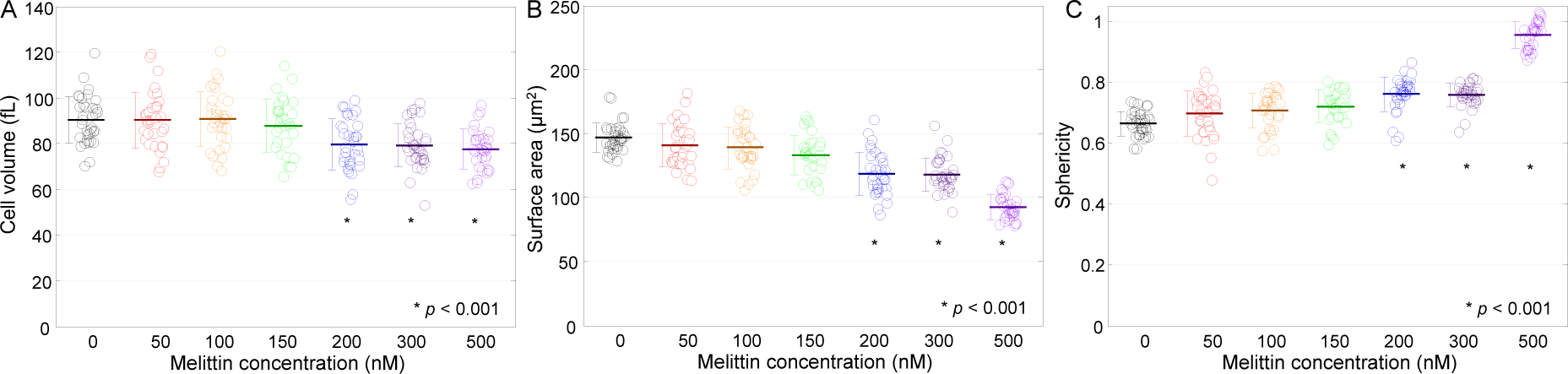
(A-C) Morphological red cell indices of RBCs at various melittin concentrations of 0, 50, 100, 150, 200, 300, and 500 nM: (A) Cell volumes, (B) surface areas, and (C) sphericities. Each open circle corresponds to an individual RBC measurement. The solid horizontal line is the mean value with an error bar for the standard deviation.

The surface areas of the RBCs decreased as the melittin concentration increased. The mean values of the surface areas for the corresponding melittin concentrations stated above were 146.8 ± 11.3, 140.7 ± 16.9, 138.9 ± 16.2, 133.1 ± 15.2, 118.3 ± 17.0, 117.8 ± 12.8, and 92.4 ± 9.8 μm^2^, respectively. From the measured cell volumes and surface areas, the sphericities were calculated. The sphericities were almost unchanged up to a concentration of 150 nM. At concentrations of 200 and 300 nM, the sphericities increased from 9% to 13.5%. At a concentration of 500 nM, the sphericity became close to unity indicating spherocytosis. The mean values of sphericity for melittin concentrations of 0, 50, 100, 150, 200, 300 and 500 nM were 0.665 ± 0.040, 0.696 ± 0.075, 0.706 ± 0.059, 0.721 ± 0.053, 0.761 ± 0.057, 0.759 ± 0.037, and 0.955 ± 0.045, respectively.

### Melittin-induced biochemical changes in RBCs

The RI map inside a cell reflects information about the constituents, distributions, and concentrations of cytoplasmic substances, and it has been used in several studies on biological objects including RBCs^24^, plankton^26^ and lipid droplets^37^. Because the cytoplasm of RBCs consists mainly of Hb, the cytoplasmic Hb concentration of individual RBCs can be calculated from the measured RI because the RI of the Hb solution is directly related to the concentration of Hb^38,39^. Furthermore, the cellular dry mass, dominantly from cytoplasmic Hb of RBCs, is accurately measured from 2-D phase images^24,40-42^.

As shown in Fig. 1, the intracellular RI value increased as the applied melittin concentration was increased, which indicates an increase in the Hb concentration. To quantitatively analyse this effect, measured Hb concentrations and dry masses of RBCs were extracted from 3-D RI maps and 2-D phase images, respectively (Fig. 3). The dry masses, interestingly, presented neither statistically distinguishable changes (*p*-value > 0.1) nor any conceivable trends despite the increasing melittin concentrations, which suggests that Hb in RBCs did not significantly leak due to melittin at nanomolar concentrations. The mean values of dry mass were 31.1 ± 3.3, 30.5 ± 3.5, 31.9 ± 3.1, 31.5 ± 3.9, 31.3 ± 3.6, 32.3 ± 3.7, and 30.6 ± 3.5 pg for melittin concentrations of 0, 50, 100, 150, 200, 300, and 500 nM, respectively. Hence, the Hb concentrations of the RBCs for melittin concentrations between 150 and 200 nM were expected to increase in response to the decrease in cell volume, and the measured Hb concentrations of the RBCs agreed with this expectation. The mean values of intracellular Hb concentration were 34.5 ± 2.8, 33.7 ± 4.7, 35.5 ± 4.9, 34.3 ± 4.1, 43.7 ± 7.4, 43.9 ± 6.2, and 49.1 ± 5.0 g/dL for the corresponding melittin concentrations, showing step-wise change between 150 and 200 nM.

**Fig. 3.**
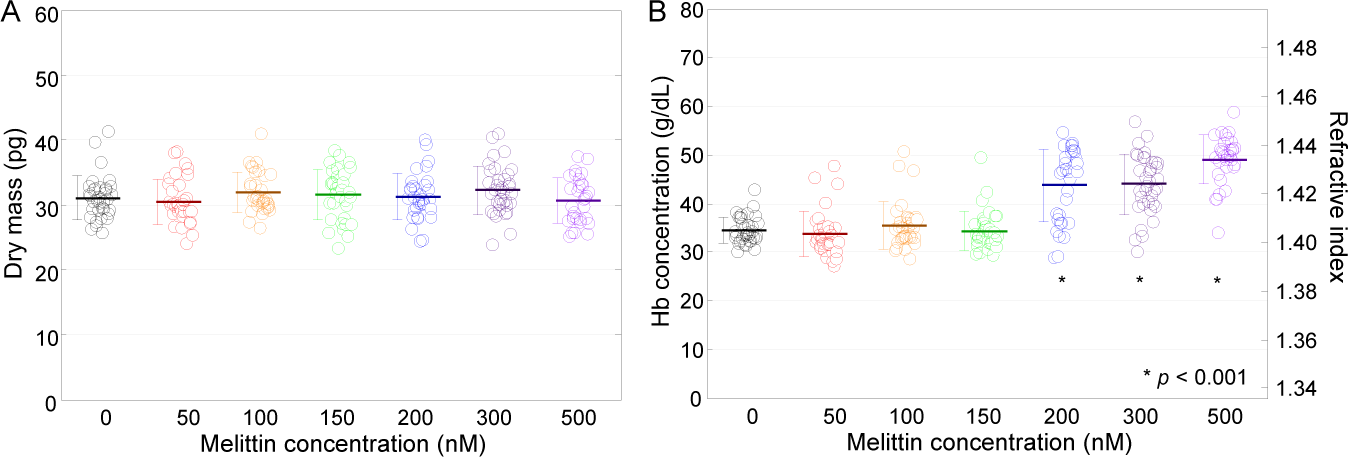
Dry mass (A) and Hb concentration (B) of RBCs at various melittin concentrations of 0, 50, 100, 150, 200, 300, and 500 nM.

### Mellitin-induced stiffening of RBC membranes

Dynamic fluctuation of the cell membrane manifests deformability of the cell membrane, which is crucial to understand the mechanical properties of the RBC due to the underlying cytoskeletal network^29,30,33,34^. Here, melittin-induced changes in the deformability of RBCs were investigated as a function of melittin concentration. From the measured 2-D phase delay maps of RBCs and intracellular RI from 3-D tomograms, the height maps of individual RBCs were retrieved (Figs. 4A–D). Height maps exhibited significant morphological changes of the same characteristics with the results from 3-D isosurface images shown in Fig. 1.

**Fig. 4.**
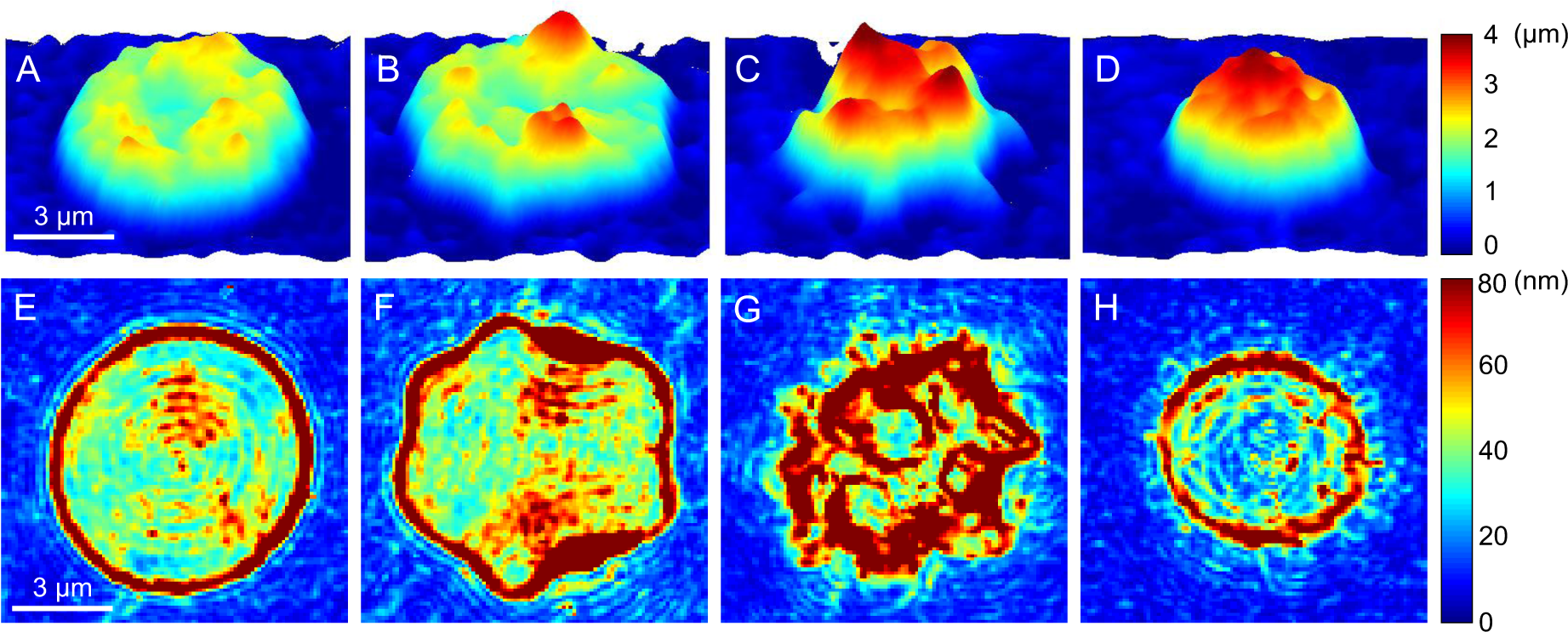
(A-D) Representative 2-D topographical images of RBCs at various melittin concentrations of 0, 50, 200, and 500 nM, respectively. (E-H) Corresponding membrane fluctuation displacement maps.

In order to quantify the deformability of individual RBCs, the membrane fluctuations were calculated from root-mean-squared *(RMS)* height displacement; the measurement of phase delay maps via common-path holographic imaging technique allows to obtain membrane fluctuation maps with nanometer precision^27–29^. The representative membrane fluctuations of single RBCs at the melittin concentrations of 0, 50, 200 and 500 nM, are presented in Figs. 4E–H, respectively. The *RMS* height displacements of individual RBCs also showed significant changes in fluctuation profiles: discocytes and spherocytes exhibited homogeneous fluctuation at intracellular regions (Figs. 4E, H), whereas echinocyte presented inhomogeneous fluctuation (Figs. 4F, G).

The mean membrane fluctuation of RBCs at each melittin concentration was obtained by the spatially averaged *RMS* height displacement fluctuation over the cell surface (Fig. 5). The mean values of fluctuation were 42.3 ± 4.7, 48.7 ± 5.8, 46.8 ± 6.4, 48.2 ± 5.5, 47.0 ± 5.7, 50.1 ± 8.7, and 34.5 ± 7.3 nm at melittin concentrations of 0, 50, 100, 150, 200, 300 and 500 nM, respectively. The fluctuation level decreased by 18% at 500 nM compared with that without melittin. The reduced fluctuation level of the spherocytes at 500 nM suggests a loss of deformability of the RBCs. This result agrees with a previous report that spherocytes and echinocytes induced by ATP-depletion exhibited suppressed fluctuation due to higher bending, shear, and area moduli than that of discocytes^30^. However, the mean fluctuation of echinocytes at 50-300 nM was measured to be higher than that of discocytes. Note that the high fluctuation signals around the spikes of the echinocytes (Figs. 4C–D) from the radial motion of the spikes rather than the normal fluctuation of the cellular membrane can be attributed to this apparent increase in the measured fluctuation level.

**Fig. 5.**
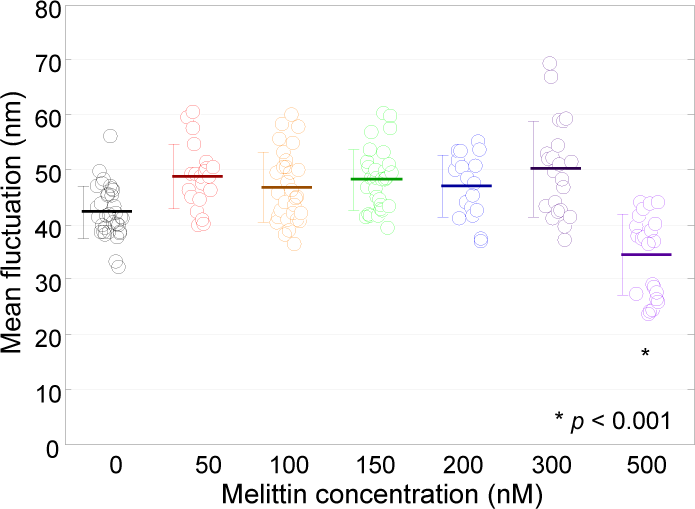
Mean membrane fluctuations of individual RBCs at various melittin concentrations of 0, 50, 100, 150, 200, 300, and 500 nM.

### Real-time measurements of Hb leakage from RBCs induced by melittin

To investigate the process of melittin-induced cellular changes as well as Hb leakage at micromolar melittin concentration, we measured the changes in a single RBC after treated with melittin via the realtime cDOT. Recent development of 3-D optical tomography allows rapid measurement of 3-D dynamics of microscopic objects at sub-second time resolution (28) which is sufficient to observe the process of melittin-induced cellular changes over time. To achieve real-time 3-D tomograms, the total number of illumination angles was reduced as 15 so that the total measurement time for each tomogram was 30 msec.

Three dimensional isosurface images clearly visualized time-dependent morphological and biochemical changes in RBC (Fig. 6). The morphological changes of individual RBCs due to melittin over time also followed the process of echinocytosis: from discocyte to echinocyte (Figs. 6A–C), and to spherocyte (Figs. 6C–D). In addition, RI inside RBCs increased as the time increased when melittin concentrations were 250 nM, 1 μM, and 5 μM before Hb started to leak. For the case of 5 μM, the rapid decrease of RI inside the cell was observed which indicates the leakage of Hb as shown in Fig. 6D. The final morphologies and increase of RI at lower melittin concentrations agreed with previous results (Fig. 1).

**Fig. 6.**
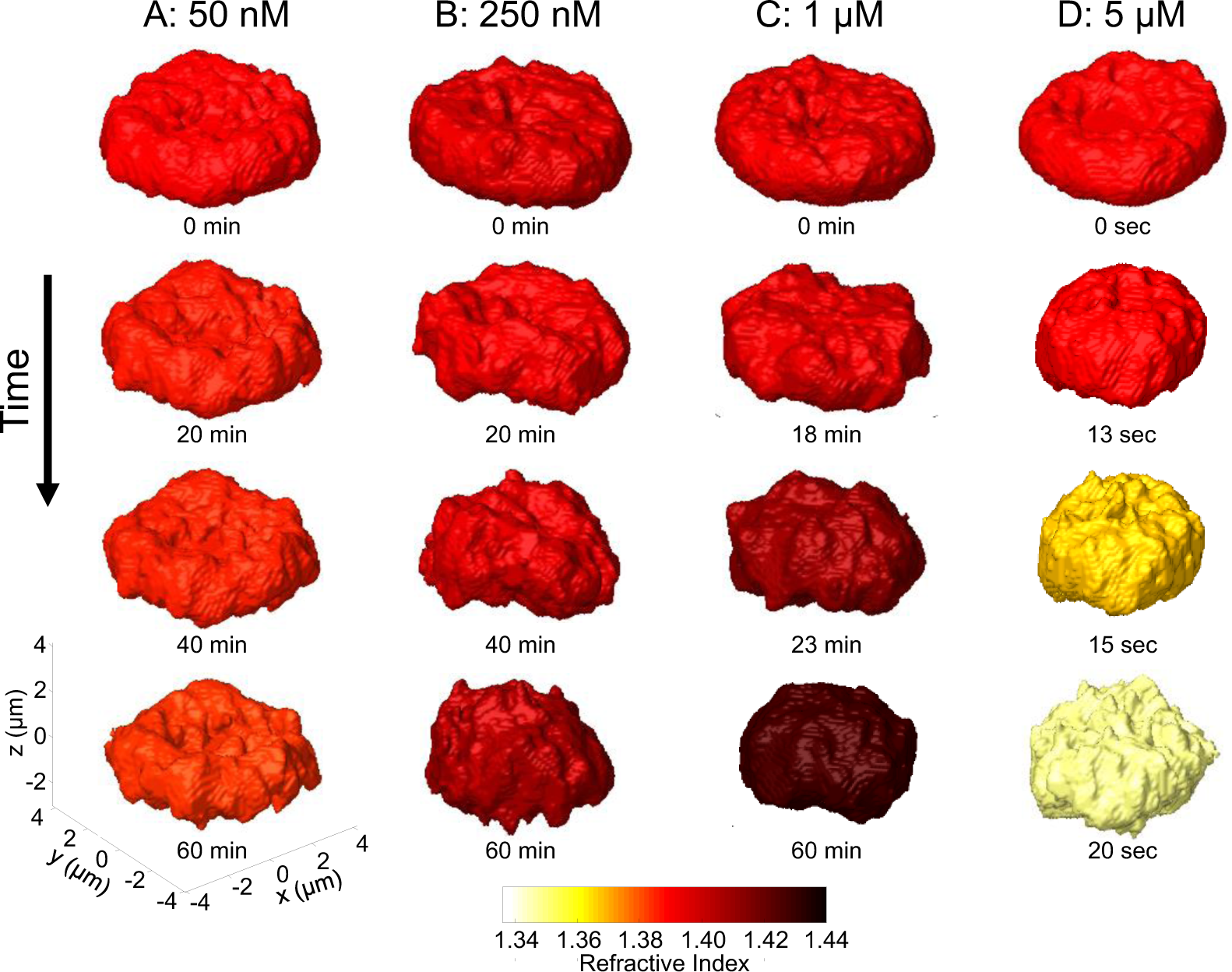
Rendered RI isosurface images of RBCs change over time at various melittin concentrations: (A) 50 nM, (B) 250 nM, (C) 1 μM, and (D) 5 μM.

The temporal changes of morphological and biochemical parameters were investigated further. The results showed the same tendencies with those of statistical studies (Figs. 2–3): as shown in Fig. 7, cell volumes and surface areas of RBCs at melittin concentrations of 50 nM, 250 nM and 1 μM decreased as time increased (Figs. 7A–B). Sphericities and Hb concentrations increased, whereas dry masses were maintained (Figs. 7D–E).

**Fig. 7.**
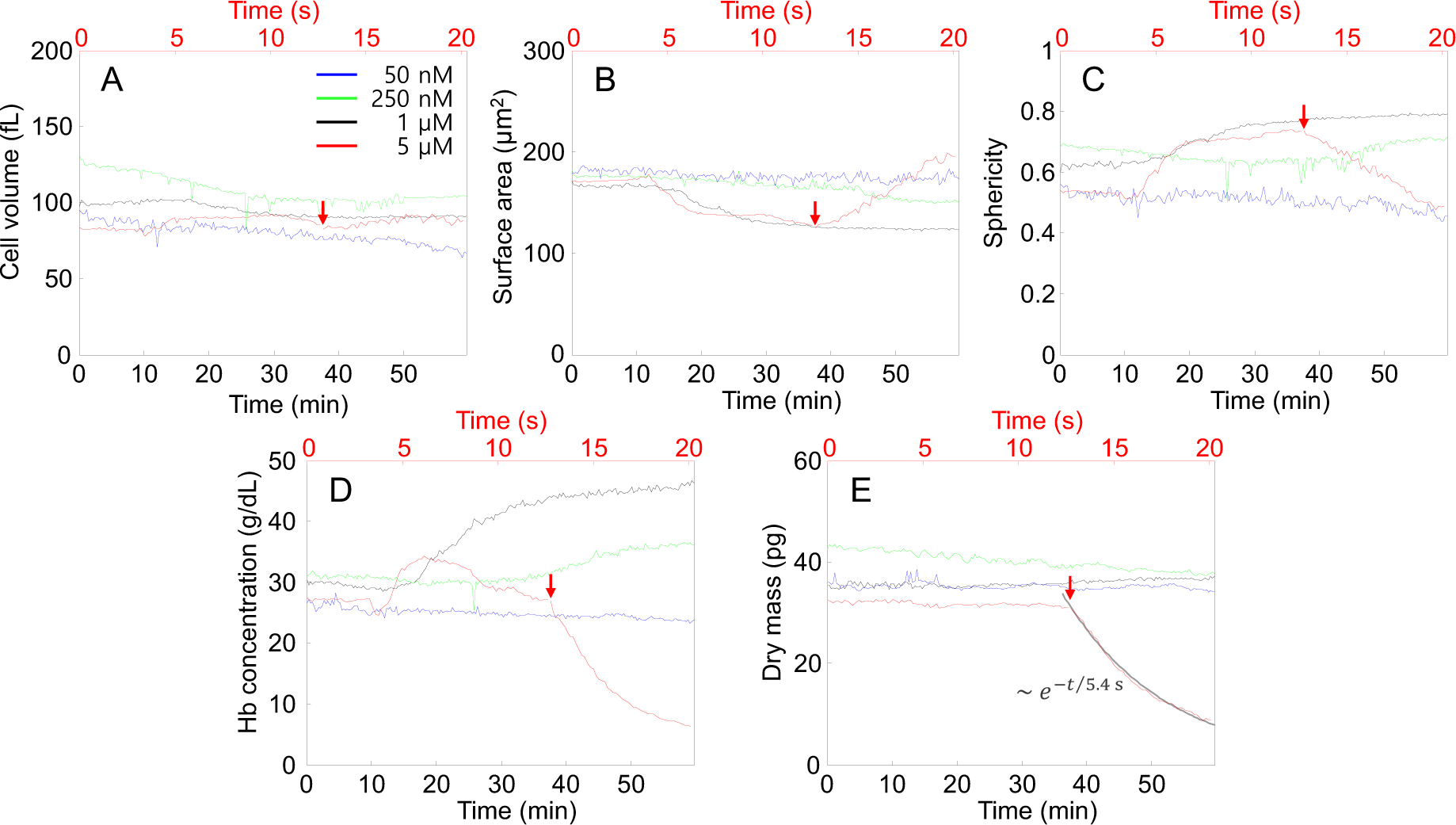
Morphological and biochemical changes of RBCs over time when melittin solutions of 50 nM (blue lines), 250 nM (green lines), 1 μM (black lines), and 5 μM (red lines) were added: (A) cell volumes, (B) surface areas, (C) sphericities, (D) Hb concentrations, and (E) dry masses. The bottom *x*-axis with the black colour is for the time elapsed in one minute for the 50 nM, 250 nM, and 1 μM cases. The top x-axis with the red colour is for the 5 μM case in one second. The red arrows indicate the start of Hb leakage. The grey line in E is the fitted exponential decay.

The variations in detailed aspects and ranges of cellular changes in the real-time measurements and statistical study were presumably due to cell-to-cell variations in the original conditions and the susceptibility of cells to the action of melittin. It is also noteworthy that the melittin-induced changes of RBCs were established within one hour after the melittin solutions were added, for melittin concentrations up to 1 μM.

When the concentration of melittin was 5 μM, fast leakage of Hb was observed 13 sec after the addition of melittin. As shown in Fig. 7D–E, the dry mass and Hb concentration underwent exponential decays. The characteristic time for the exponential decay of the dry mass was 5.417 sec. While the Hb concentration and dry mass decreased, the cell volume did not change significantly, and the surface area increased. The increase in the surface area is explained as follows: as the Hb concentration decreased by the Hb leakage, the 3-D isosurface images became sensitive to the noise of the RI map and presented more noisy surfaces shown in Fig. 6D, which resulted in the increase in the surface area and sphericity due to their definitions. The inhomogeneity of the Hb concentration during Hb leakage also contributed to the formation of noisy isosurfaces.

## Discussion and Conclusion

In this study, melittin-induced morphological, biochemical and mechanical changes in RBCs were investigated with real-time measurements from 3-D RI tomograms and 2-D dynamic phase maps using cDOT. Morphological changes of RBCs from discocytes to echinocytes or spherocytes by binding of melittin, a pore-forming protein, were observed. In addition, the morphologies were dependent on the concentrations of melittin. Geometrical parameters were extracted from the 3-D RI tomograms; cell volumes and surface areas exhibited decreases as the melittin concentration increased while the sphericities increased indicating the progression of echinocytosis and spherocytosis. The biochemical conditions were measured from the intracellular RIs: the dry masses of the RBCs did not change as melittin solutions of increasing concentrations were added, whereas the Hb concentrations did increase. Membrane fluctuation profiles showed that the mean membrane fluctuation levels decreased when the RBCs became spherocytes from the melittin treatment suggesting that melittin reduced the deformability of the RBCs. The measured values of the cell volume, surface area, sphericity, Hb concentration, dry mass, and mean fluctuation level of the control (0 nM) were in good agreement with previous results^30,43-46^. The real-time measurements showed the characteristic time scale of the melittin-induced changes and enabled us to study the Hb leakage process at high melittin concentrations. The dry mass of an RBC underwent exponential decay during Hb leakage, and the characteristic time of the decay was fitted for further analysis.

The effect of melittin to induce echinocytosis and spherocytosis can be explained by the fact that melittin is a strong stimulator of phospholipase a2 (PLA2)^47^. Previous reports showed that PLA2 in snake venom is responsible for echinocytosis of RBCs by producing lysolecithin, a known echinocytic agent^48–50^. The leakage of adenosine triphosphate (ATP) through melittin-induced stable pores could accelerate echinocytosis because the depletion of cytoplasmic ATP is also known as a trigger of echinocytosis or spherocytosis.

During morphological transformation by melittin binding, stiffening of the spectrin network beneath the RBC membrane occurs and results in the loss of the membrane through vesiculation and a decrease in the surface area. Microscopic vesiculation under echinocytic condition was suggested in previous studies by Park *et al.*^30^ and Sens and Gov^51^. Sens and Gov proposed that the vesiculation of RBCs is the result of stiffening of the underlying spectrin network, which can also explain our result that the membranes of the RBCs lost their deformability by melittin binding. Furthermore, the stiffening of the spectrin network could be from not only microscopic vesiculation but also from changes in the global geometry of the entire cell membrane. Therefore, a step-wise decrease in the cell volume (Fig. 2A) is expected to be related to a change in the global shape of the RBC, and we suggest that the loss of the dimples in the centres of RBCs is correlated with the decrease in the cell volume by analysing the curvatures of individual RBCs (See supplementary information 1).

With micromolar concentrations of melittin, a decrease in the intracellular Hb concentration over time was observed without significant changes in the cell volume (Figs. 6D and 7). Considering the pore-forming effect of melittin^1^ and the exponential decay of dry mass as well as Hb concentration, the leakage of Hb should be outward diffusion of intracellular Hb through melittin-induced pores in the membrane because the diffusion current is proportional to the concentration contrast between the cells and medium. Applying the diffusion layer model

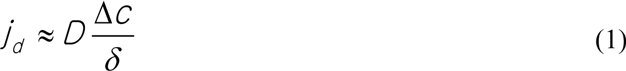

where *j*_*d*_ is the current density of diffusion; *D* is the diffusion coefficient; Δc is the Hb mass concentration contrast between the cytoplasm and medium, and *δ* is the effective thickness of a diffusion layer (i.e., membrane of RBC), we could establish a rough estimation as follows:

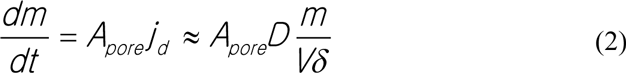

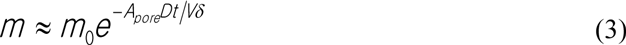

with a characteristic time of *τ* ≈ *Vδ/A*_*pore*_*D,* where *m* is the mass of Hb inside the cell, and A_pore_ is the total area of the pores in the cell membrane. By applying *D* = 70 μm^2^/s for Hb^52^, δ = 7.8 nm as the thickness of the RBC membrane^53^, *V* = 82.6 fL as the measured cell volume at the start of Hb leakage (Fig. 7A), and τ = 5.4 s as the measured characteristic time of the exponential dry mass decay (Fig. 7E), A_pore_ was estimated as 1700 nm^2^. Assuming the cylindrical pores have a uniform diameter at 5 nm and are constant over time and comparable to the diameters of the Hb molecules^1,54^, the number of pores in the membrane was estimated as 87. Repeated measurements of the Hb leakage led to statistical results (See supplementary information 2): the characteristic times were 4.85 ± 1.58 and 1.19 ± 0.49 s; the estimated total areas of the pores were 2061 ± 632 and 8675 ± 2890 nm^2^, and the numbers of pores were 105 ± 32 and 442 ± 147 for melittin concentrations of 5 and 10 μM, respectively.

This study reports for the first time the 3-D investigation of the effects of melittin on RBCs in terms of both the biophysical alterations of RBCs at low melittin concentrations and haemolysis at high concentrations. There have been many studies on transmembrane proteins related to human health including Gp41 in HIV^55^, haemolysins^56^, and melittin^1^ for the various therapeutics including medications, antibiotics, and selective attack. Furthermore, studies on designing and synthesizing artificial transmembrane channels have been conducted for three decades^57–61^. However, most of these studies concentrated on microscopic effects (molecular binding on the membrane, pore formation, functions of the channel, etc.), even though the effects of those proteins on live cells on a whole cell scale must also be examined before clinical use. Therefore, we suggest that the systematic approach using cDOT in this study is suitable to study the cellular effects of other transmembrane proteins or, even more general, to study the effects of any biochemical on various cells including white blood cells and cancer cells.

## Methods

### Sample preparation (Ethics statement) IRB

Human blood studies were done according to the principles of the Declaration of Helsinki and approved by the institutional review board of KAIST (IRB project number: KH2013-22, Daejeon, Republic of Korea). All experimental protocols were approved by the institutional review board of KAIST. The participant provided written informed consent to participate in this study which was approved by the ethics committees and IRB.

A 3 μL drop of blood from a fingertip was collected from a healthy donor and immediately diluted 37.5-fold in Dulbecco’s Phosphate Buffered Saline (PBS) solution (Gibco®, New York, U.S.A.). The melittin protein (M2272, Sigma-Aldrich), purified from honey bee venom, was also prepared in PBS solution. The diluted blood was added to the melittin solution for various desired melittin concentrations, and the final dilution factor of the blood was kept at 300-fold. Each RBC with melittin sample was prepared at room temperature.

### Common-path Diffraction Optical Tomography (cDOT)

The details of the cDOT setup were described in a previous work^32^. A diode-pumped solid-state laser (λ= 532 nm) was used as an illumination source for an inverted microscope. At the sample plane, the illumination angle of the beam was rapidly scanned by rotating the first galvanometric mirror, which was placed before the sample stage. The second galvanometric mirror, which was located after the sample stage, rotated in the opposite direction of the first mirror such that the beam reflected from the second mirror had the same optical path regardless of the illumination angle. The optical field of the diffracted beam was quantitatively and precisely measured with a common-path interferometry setup: the beam from a sample was diffracted by a grating. The 0^th^ order diffraction beam was spatially filtered by a pinhole with a diameter of 25 μm to serve as a reference plane wave at the image plane. The 1^st^ order beam was directly projected onto the image plane. At the image plane, the sample and reference beams interfered forming spatially-modulated interferograms. Interferograms were recorded by a CCD camera (Neo sCMOS, ANDOR Inc., Northern Ireland, UK) with a pixel size of 6.5 μm and x 240 total magnification of the imaging system, and the optical field with both amplitude and phase information was quantitatively retrieved with a phase retrieval algorithm^62^. By changing the angles of illumination impinging on the sample, cDOT measures multiple 2D optical fields with different illumination angles from which the 3-D RI tomogram of the sample is reconstructed with the diffraction tomogram algorithm^31,63,64^. The 2-D dynamic height profiles were measured at the fixed normal illumination angle.

### RI tomogram measurement and red cell parameter analysis of melittin bound RBCs

RBCs samples with melittin concentrations of 50, 100, 150, 200, 300, and 500 nM were prepared. Then, we waited one hour after the melittin solutions were mixed with the RBCs. Three-dimensional RI tomograms and 2-D dynamic phase images of the RBCs were measured at the single-cell level. The appropriate upper bound was 500 nM because the RBCs started to exhibit a significantly lower RI and smaller size at a micromolar concentration (See Mov. 4). RBCs without melittin (0 nM) were measured as the control. A droplet of each sample solution was sandwiched between two cover glasses (C024501, Matsunami Class Ind., Japan) spaced apart by melted parafilm strips for the cDOT measurement.

For each 3-D tomogram measurement of a single cell, a set of 300 interferograms with various angles of illumination was recorded at a frame rate of 100 Hz. Isosurface images of 3-D RI tomograms with a threshold RI value of 1.368 were taken to measure the cell volumes and surface areas. The sphericities of individual RBCs defined by (36*πV*^2^)^1/3^/*A* where *V*is the volume and *A* is the surface area of a cell, were calculated from the measured cell volumes and surface areas. The mean RI values inside individual cells were translated to the Hb concentrations of the RBCs using a model function^39^. The 2D phase delay maps of the RBCs were measured at the fixed normal illumination angle at a frame rate of 125 Hz for 2 sec. The cell height profiles, *h(x,y;t),* of the RBCs were retrieved from the measured phase delay map, Δ_*φ*_(*x,y;t*), using the following relation:

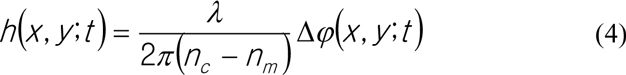

where is the wavelength of the laser light, and n_c_ and *n*_*m*_ are the RI of the cell cytoplasm and surrounding medium, respectively. The fluctuation map Δ*h*(*x*,*y*) was obtained by taking the root-mean-square of the height profiles over time at each position. The mean fluctuation of the individual RBCs was obtained as the mean values of the local fluctuation Δ*h*(x,y) inside the cells. The dry mass of individual RBCs, which mainly consist of cytoplasmic Hb, was measured from the 2-D phase delay maps with the following relationships^42^:

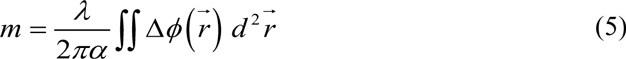

where *α* is the RI increment of the Hb solution^39^.

### Real-time measurement of melittin-induced changes in RBCs

Two cover glasses, one shorter than the other (W × L = 24 × 50 mm and 24 × 40 mm, C024501 and C024401, Matsunami Class Ind., Japan), were spaced apart with a pair of melted parafilm strips to form a narrow channel. To adhere the cell membrane of the RBCs to the surface of the bottom glass, the channel was treated with 0.000005% poly-L-lysine solution (P8920, Sigma-Aldrich) for 15 minutes, then washed 3 times with PBS buffer, and finally placed on the sample plane in the cDOT setup. The dilute RBC solution was dropped onto one side and introduced into the channel by capillary action. The amount of the RBC solution was adjusted so that only half of the channel was occupied. After that, the melittin solution with the desired concentration was dropped onto the same edge so that the channel was completely filled. For each melittin concentration of 50 nM, 250 nM, and 1 μM, 15 interferograms of individual RBCs with various angles of illumination were measured at a frame rate of 100 Hz, and in total, 200 sets of interferograms were recorded at a period of 18 sec. for one hour. For the 5 μM case, 200 sets of interferograms were recorded at a period of 150 ms for 30 sec.

## AUTHOR CONTRIBUTIONS

J.H., H.P., and Y.P. developed the experimental idea. J.H. performed the sample preparation and optical measurement. K.K. and S.L. built the optical setup and established the image processing algorithm. J.H. analyzed the data. Y.P. and H.P. conceived and supervised the study. All authors discussed the experimental results and wrote the manuscript.

## ACKNOWLEDGEMENTS

This work was supported by KAIST, Tomocube, and the National Research Foundation of Korea (2015R1A3A2066550, 2014K1A3A1A09063027, 2014M3C1A3052567) and Innopolis foundation (A2015DD126).

